## Supplementary material for "Joint copy number and mutation phylogeny reconstruction from single-cell amplicon sequencing data"

#### Contents

**A Details on the probabilistic model** **2**

**B MCMC moves for exploring the tree space** **4**

**C Properties of the Tapestri data** **10**

**D Simulations with synthetic data** **16**

**E Orthogonal validation of COMPASS-derived CNA calls with bulk data** **20**

**F Additional trees** **26**

### A Details on the probabilistic model

#### A.1 Notations

Cells are indexed by the letter $j$, variants by $i$, regions by $k$ and nodes by $n$.

- $D_{ij}$: sequencing depth at locus $i$ for cell $j$ (and $D_{kj}$ is the depth at region $k$ for cell $j$, and $D_j$ is the total number of reads in cell $j$).
- $A_{ij}$: number of alternative reads at locus $i$ for cell $j$.
- $\sigma_j$: node to which cell $j$ is attached.
- $\pi_n$: prior probability of sampling a cell from node $n$.
- $c_k(n)$: copy number of region $k$ for node $n$.
- $c_i^{(r)}(n)$: copy number of the reference allele at locus $i$ for node $n$.
- $c_i^{(a)}(n)$: copy number of the alternative allele at locus $i$ for node $n$.
- $\mu_i$: dropout rate for locus $i$. When separate dropout rates are used for the two alleles, $\mu_i^{(r)}$ and $\mu_i^{(a)}$ indicate respectively the dropout rate of the reference and alternative allele.
- $\lambda$: probability that the two alleles for one locus have a different dropout rate, which is constant and was set to $e^{-70}$.
- $\alpha, \beta$: parameters for the Beta distribution corresponding to the prior on dropout rates, which are constant and were set to 5 and 95 respectively.
- $C_{ij}^{(r)}$: Number of copies of the reference allele for locus $i$ that did not get dropped out in cell $j$.
- $C_{ij}^{(a)}$: Number of copies of the alternative allele for locus $i$ that did not get dropped out in cell $j$.
- $\varepsilon$: sequencing error rate (the same is used for all sites), which is constant and was set to 1%.
- $\omega_{\text{hom}}$: overdispersion of the beta binomial when the cell is homozygous for a locus (a high $\omega$ corresponds to a low variance), which is constant and was set to 50.
- $\omega_{\text{het}}$: overdispersion of the beta binomial when the cell is heterozygous for a locus, which is constant and was set to 8.
- $\delta$: doublet rate, which is constant and was set to 8%.
- $\rho_k$: probability for a read to fall on region $k$, when there are two copies of this region.
- $\theta$: inverse dispersion parameter of the negative binomial distribution, which is constant and was set to 6.

#### A.2 Generative process

Given a mutation tree, we assume the following generative process for the read counts, which is used to derive likelihoods.

- For each variant $i$:

  - With probability $\lambda$, draw the dropout rate of both alleles $\mu_i^{(r)}$ and $\mu_i^{(a)}$ independently from a Beta distribution with parameters $\alpha$ and $\beta$ (we used $\alpha = 5$ and $\beta = 95$ so that the prior distribution has a mean of 0.05).
  - With probability $1 - \lambda$, both alleles have the same dropout rate. Draw $\mu_i = \mu_i^{(r)} = \mu_i^{(a)}$ from a Beta distribution with parameters $\alpha$ and $\beta$.

- Draw the node weights $\pi_1, ...\pi_N$ from a flat Dirichlet distribution $D(1, ..., 1)$.
- For each cell $j$:
  - With probability $1 - \delta$, the cell is a singlet.
    * Draw the attachment node $\sigma_j$ from Categorical$(\pi_1, ...\pi_N)$.
    * Draw the total number of reads in the cell $D_j$.
    * For each region $k$:
      · draw the number of reads falling on region $k$ $D_{kj}$ from a negative binomial distribution $\mathrm{NB}(D_j \frac{c_k(\sigma_j)}{2} \rho_k, \theta)$.
    * For each variable locus $i$:
      · Draw the number of copies of the reference allele that are not dropped out $C_{ij}^{(r)}$ from $\mathrm{Binom}(c_i^{(r)}(\sigma_j), 1 - \mu_i^{(r)})$
      · Draw the number of copies of the alternative allele that are not dropped out $C_{ij}^{(a)}$ from $\mathrm{Binom}(c_i^{(a)}(\sigma_j), 1 - \mu_i^{(a)})$
      · Draw the alternative read counts from $\mathrm{BetaBinom}(D_{ij}, f, \omega)$ where $f = \frac{C_{ij}^{(a)}}{C_{ij}^{(r)} + C_{ij}^{(a)}}(1-\varepsilon) + \frac{C_{ij}^{(r)}}{C_{ij}^{(r)} + C_{ij}^{(a)}}\varepsilon$ is the expected frequency of the alternative allele and the overdispersion $\omega$ is $\omega_{\text{hom}}$ if $C_{ij}^{(r)} = 0$ or $C_{ij}^{(a)} = 0$ and $\omega_{\text{het}}$ otherwise.
  - With probability $\delta$, the cell is a doublet.
    * Draw the two nodes independently, each from Categorical$(\pi_1, ...\pi_N)$.
    * The genotype of the doublet is obtained by summing the copy numbers of each node.
    * Given the genotype of the doublet, the reads are generated in the same way as for a singlet.

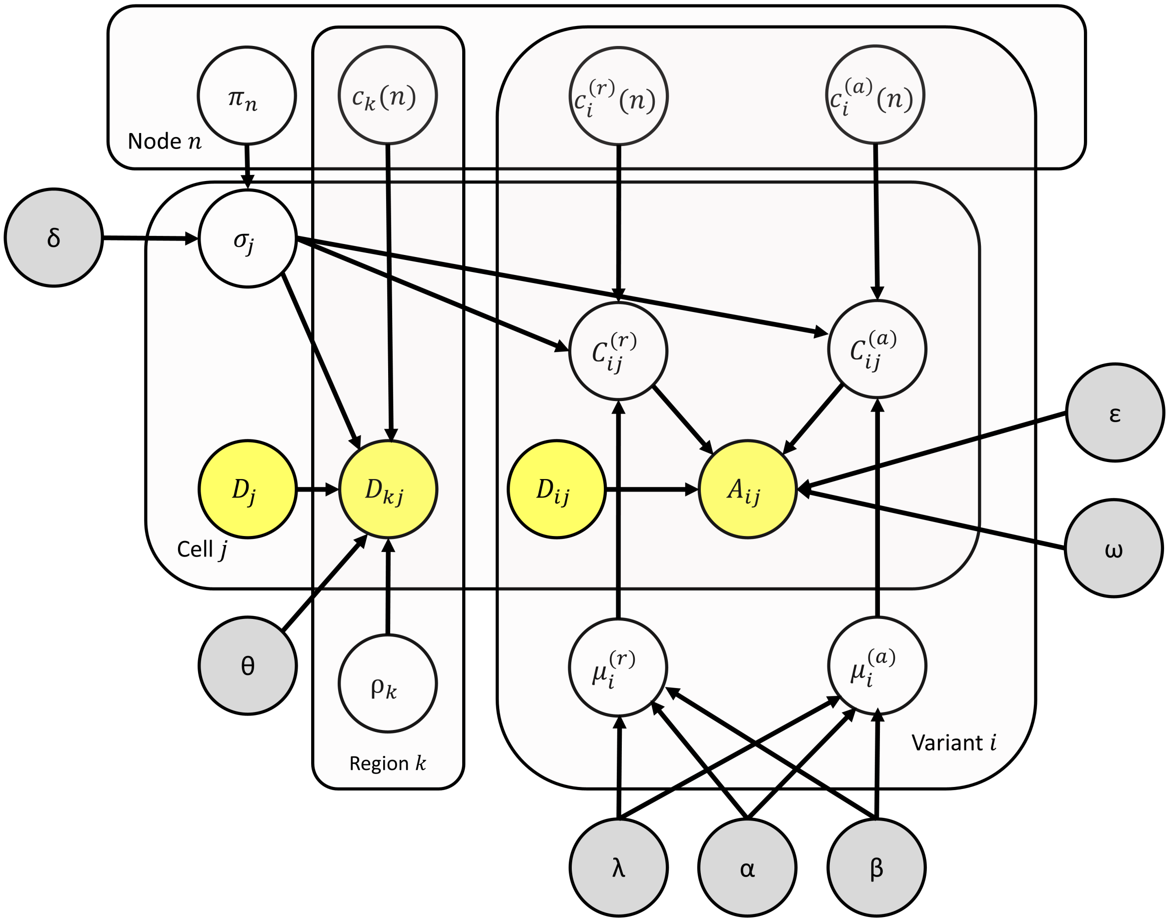


Supplementary Figure A.1: Probabilistic graphical model representing the generative process. Yellow nodes correspond to observed variables and grey nodes to constants. All of the other variables are estimated in the inference.

### B MCMC moves for exploring the tree space

In this section, we describe in more detail each move used to propose a new tree in the MCMC. Most of these moves are not symmetric (proposing $\mathcal{T}'$ from $\mathcal{T}$ does not have the same probability as proposing $\mathcal{T}$ from $\mathcal{T}'$), so we also explain how to compute the Hastings ratio.

#### B.1 Prune and reattach

- Select one node $n_1$ uniformly at random.
- Select one node $n_2$ which is not a descendant of $n_1$.
- Set $n_2$ as the new parent of $n_1$ (which results in moving the whole subtree below $n_1$).

This move is symmetric, so the Hastings ratio is 1:

$$\mathrm{HR} = 1$$

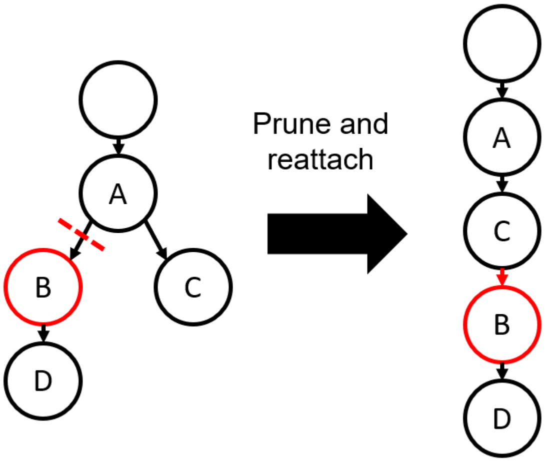


Supplementary Figure B.1: Prune and reattach a node (here node B). The subtree (here node D) is moved along with the node.

#### B.2 Swap node labels

- Select one node $n_1$ uniformly at random.
- Select one node $n_2$ uniformly at random among all other nodes.
- Exchange the events in $n_1$ with those in $n_2$.

This move is symmetric, so the Hastings ratio is 1:

$$\mathrm{HR} = 1$$

#### B.3 Move event

- Select a type of event among SNV and CNA.
- Select a source node $n_1$ which has this type of event ($N_{\text{event}}$ possibilities).
- Select one event in this node ($n_{\text{event}}(n_1)$ possibilities) and remove it.
- If the node is now empty, delete it and set the parent of its children to be the parent $p$ of the deleted node.
- Choose whether to move the event to a new node, or to an existing node.
  - With probability $\alpha$, move the event to a new node $n_2$.
    - Select a parent $p$ for the new node among all the existing nodes ($N$ possibilities)
    - For each child of the parent, reassign its parent to the new node with probability 1/2. ($2^{n_{\text{children}}(p)}$ possibilities).
  - With probability $1 - \alpha$, move the event to an existing node $n_2$.
    - Select the node among all the existing nodes ($N$ possibilities)
- If we moved an SNV and the tree contains CNAs which affect this locus, randomly change the allele affected by the CNA for this locus.

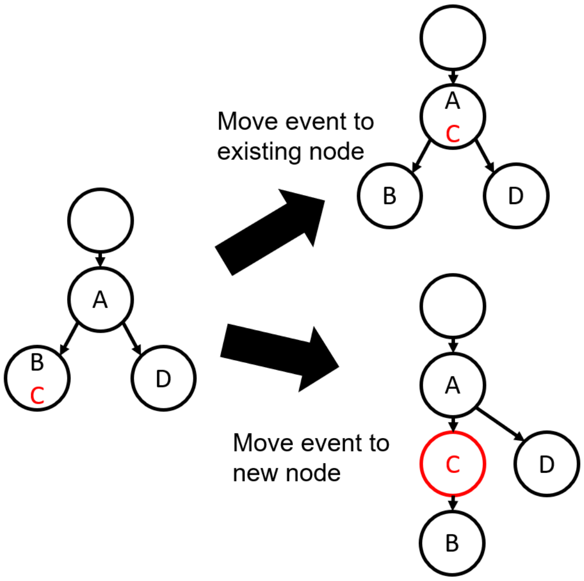


Supplementary Figure B.2: Move a somatic event. Here, event C can either be moved to an existing node (top) or to a new node (bottom). When an event is moved to a new node, the new node has to select its parent among the existing nodes, and the children are randomly reassigned: here, the node containing event D keeps A as its parent, but the node containing event B has the new node as its parent.

To reverse this move, we need to select $n_2$ as the source node ($N'_{\text{event}}$ possibilities, because there might be one more or one fewer node with this type of event, after moving the event), select the right event in $n_2$ ($n_{\text{event}}(n_2)$ possibilities). If $n_1$ was deleted, we need to select a new node as a destination node, select the right parent and reassign the children correctly. Otherwise, we need to select $n_1$ as the destination node ($N-1$ possibilities). Let $\beta = \alpha$ if we moved the event to a new node and $\beta = 1-\alpha$ otherwise, and let $\gamma = \alpha$ if the source node was deleted, and $1-\alpha$ otherwise. The Hastings ratio is:

$$\text{HR} = \frac{\gamma}{\beta} \frac{N_{\text{event}}}{N'_{\text{event}}} \frac{n_{\text{event}}(n_1)}{n_{\text{event}}(n_2)}$$

#### B.4 Split or merge node

Let $\alpha$ be the probability to split a node, and $1-\alpha$ is the probability to merge two nodes.

##### B.4.1 Split node

- Select one node $n$ uniformly at random ($N$ possibilities).
- Randomly allocate the events between the two nodes ($2^{n_{\text{events}}(n)}$ possibilities).
- For all of the children of the original node, set its parent to either one of the two new nodes ($2^{n_{\text{children}}(n)}$ possibilities).

To reverse the move, we need to select merge and select the right node to merge with its parent. Note that, when we merge two nodes, we cannot select the root so we have $N-1$ possibilities, but after splitting $N$ was increased by one, so $N-1$ after the split is equal to $N$ before.

$$\text{HR} = \frac{1-\alpha}{\alpha} 2^{n_{\text{events}}(n)} 2^{n_{\text{children}}(n)}$$

##### B.4.2 Merge nodes

- Select one node different from the root uniformly at random ($N-1$ possibilities).
- Merge the node with its parent (combine events of both nodes, and set the parents of their children to this merged node).

$$\text{HR} = \frac{\alpha}{1-\alpha} \frac{1}{2^{n_{\text{events}}(n)} 2^{n_{\text{children}}(n)}}$$

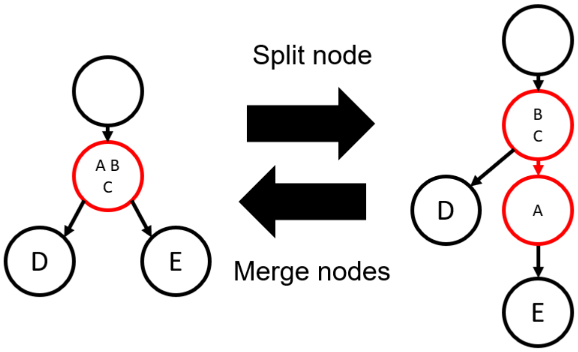


Supplementary Figure B.3: Split or merge nodes. When splitting a node, the children of the original node are randomly assigned to the two resulting nodes.

#### B.5 Add or remove CNA event

Let $\alpha$ be the probability to add a CNA event (and $1-\alpha$ the probability to remove a CNA event). $\alpha$ should be quite high (around 0.8) because there are more ways to add an event than to remove one. If there is no CNA event in the tree, we cannot remove an event so we set the probability to add an event to 1. Let $\alpha'$ be the effective add probability in this configuration, which is 1 if there is no CNA event in the tree and $\alpha$ otherwise.

##### B.5.1 Add CNA event

- Choose whether to add the event to a new node, or to an existing node.
  - With probability 1/2, add the event to a new node.
    * Select a parent $p$ for the new node among all the existing nodes ($N$ possibilities)
    * For each child of the parent, reassign its parent to the new node with probability 1/2. ($2^{n_{\text{children}}(p)}$ possibilities).
  - With probability 1/2, add the event to an existing node.
    * Select the node among all the existing nodes ($N$ possibilities)
- Select gain, loss or CNLOH (3 possibilities).

- Select a region uniformly at random . If the event is a CNLOH, can only select from the regions which contain SNVs. If the event is a gain or a loss, can only select from the candidate regions. There are $n_{\text{candidate regions}}$ possibilities.
- For each variable site in this region (if there are any), select which allele is gained or lost ($2^{n_{\text{loci in region}}}$ possibilities).

To reverse the move, we need to select remove, select this node among all nodes which have a CNA event (taking into account that, if the node initially had no CNA event, there is now one more node with a CNA event), and select the right event among all the CNA events in this node. Consequently, the Hastings ratio is:

$$\text{HR} = \frac{1-\alpha}{\alpha'} \frac{N 2^{n_{\text{children}}(p)}}{N_{\text{CNA}}} \frac{n_{\text{candidate regions}} \times 3 \times 2^{n_{\text{loci in region}}}}{n_{\text{CNA}}(n)}$$

##### B.5.2 Remove CNA event

- Select a node which has a CNA event ($N_{\text{CNA}}$ possibilities).
- Select one CNA event among all CNA events in this node ($n_{\text{CNA}}(n)$ possibilities).
- If the node is now empty, delete it and set the parent of its children to be the parent $p$ of the deleted node.

$$\text{HR} = \frac{\alpha}{1-\alpha} \frac{N_{\text{CNA}}}{N 2^{n_{\text{children}}(p)}} \frac{n_{\text{CNA}}(n)}{n_{\text{candidate regions}} \times 3 \times 2^{n_{\text{loci in region}}}}$$

#### B.6 Exchange Loss and CNLOH

- Select one node which has at least one Copy Neutral LOH event or a Copy Number Loss of a region containing at least one variant.
- Select one of such events in the node.
- If we selected a CNLOH event:
  - Remove the CNLOH event.
  - Add a Copy Number Loss of the region, with the alleles lost corresponding to the lost alleles of the original CNLOH event.
- If we selected a loss:
  - Remove the CNA event.
  - Add a CNLOH event, with the same alleles lost as in the previous loss.

The move is symmetric.

$$\text{HR} = 1$$

#### B.7 Change alleles affected by a CNA event

- Select one node containing at least one CNA event of a region containing variants.
- Select one such CNA event in that node.
- For each variant in the region affected by the CNA, randomly select which allele is affected.

This move is symmetric.

$$\mathrm{HR} = 1$$

#### B.8 Merge or duplicate CNA

The goal of this move is to allow a transition in one-step from having two identical CNAs in two different lineages to having only one copy of this CNA at their most recent common ancestor, and vice-versa.

- Count the number of CNA events which exist in several copies in the tree $n_{\mathrm{dup\_CNA}}$ and the number of nodes containing at least one CNA event and having multiple children $n_{\mathrm{nodes\_CNA\_children}}$.
- With probability $\frac{n_{\mathrm{dup\_CNA}}}{n_{\mathrm{dup\_CNA}}+n_{\mathrm{nodes\_CNA\_children}}}$, merge two identical CNA events to their most recent common ancestor.
  - Select the CNA among the $n_{\mathrm{dup\_CNA}}$ possibilities.
  - Select two nodes having this CNA ($\binom{n_{\mathrm{nodesCNA}}}{2}$ possibilities).
  - Remove this CNA from the two selected nodes, and add the CNA to their most recent common ancestor.
- With probability $\frac{n_{\mathrm{nodes\_CNA\_children}}}{n_{\mathrm{dup\_CNA}}+n_{\mathrm{nodes\_CNA\_children}}}$, duplicate a CNA event and move the two copies to two nodes in two different branches.
  - Select one node having multiple children and containing at least one CNA event ($n_{\mathrm{nodes\_CNA\_children}}$ possibilities).
  - Select one CNA event in this node and remove it.
  - Select two children of this node.
  - For each child, select one node in its subtree, and add the CNA event to it.

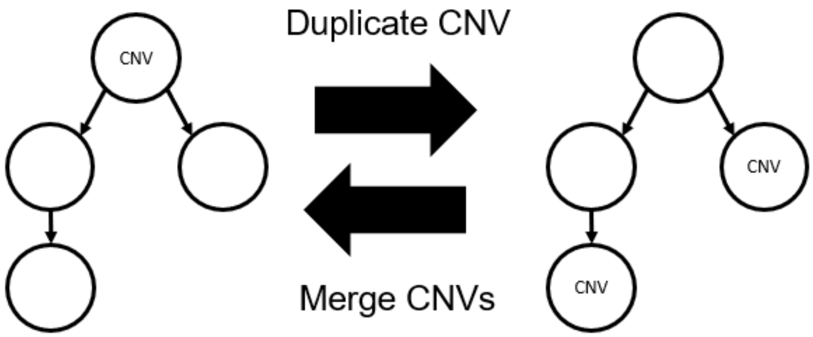


Supplementary Figure B.4: Merge or duplicate a CNA event. When merging, the two events are merged at their most recent common ancestor. When duplicating, the two copies can be placed anywhere in two distinct lineages below the original CNA.

### C Properties of the Tapestri data

#### C.1 Amplicons coverage

In Tapestri® data, each amplicon has a different amplification efficiency, with some amplicons having very low coverage to the point where most cells cannot be genotyped. Supplementary Figure C.1 shows the relationship between the GC-content of an amplicon and the percentage of reads falling on this amplicon, and we do not observe any correlation: even amplicons having a GC-content of around 50% can have a very low coverage. While an extreme GC-content might have a large impact on the coverage of an amplicon, here all the amplicons in both panels had a moderate GC-content between 30% and 70%, which is probably a consequence of having avoided amplicons with extreme GC-content in the panel design. Supplementary Figure C.2 shows for each amplicon a violin plot representing the distribution of the proportion of reads falling on that amplicon across cells, in one sample. In addition to the variability between amplicons, there is a substantial variance across cells for the same amplicon. The relative coverage of the different regions is fairly consistent across samples sequenced with the sample amplicon panel (Supplementary Figure C.3).

When there are no CNAs, we would expect the sequencing depth on each amplicon to be independent. However, this is not what is observed in practice: Supplementary Figure C.4A shows the correlation coefficients between the read counts on each amplicon in one sample, and there are very strong correlations. The same correlations are present in every sample sequenced with the 50-amplicon panel. There are also strong correlations in the 279-amplicon panel, but they have a different pattern (Supplementary Figure C.4B). With this larger panel, amplicons on the same gene tend to be correlated, but this is not always the case, and there are also numerous strong correlations between amplicons on different genes, and even between amplicons on different chromosomes (Supplementary Figure C.4C and D).

The biological explanation for these correlations is not clear. It might be that the efficiency of the proteases used to remove nuclear proteins is variable, and when they do not work well in some cells, we find a much lower sequencing depth in these cells for amplicons located in regions of tightly packaged chromatin. Such correlations in Tapestri® data have never been reported, and they have the potential to strongly confound the CNA inference, since we could interpret the two main clusters as two different clones with very different copy number profiles. However, these correlations are independent from the actual clonal architecture of the tumour, so by jointly inferring SNVs and CNAs, only the true CNAs should be detected.

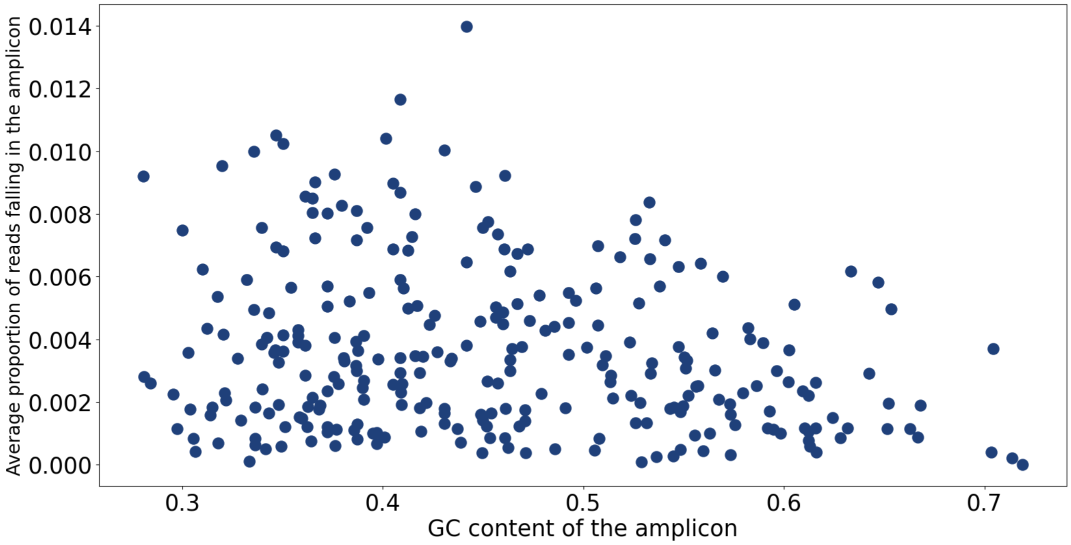


Supplementary Figure C.1: Average proportion of reads falling on each amplicon, depending on the GC-content of that amplicon, for the sample AML-72-001 (279-amplicon panel). There is a large coverage variability across amplicons, but no clear correlation with the GC-content. The situation is similar for the 50-amplicon panel.

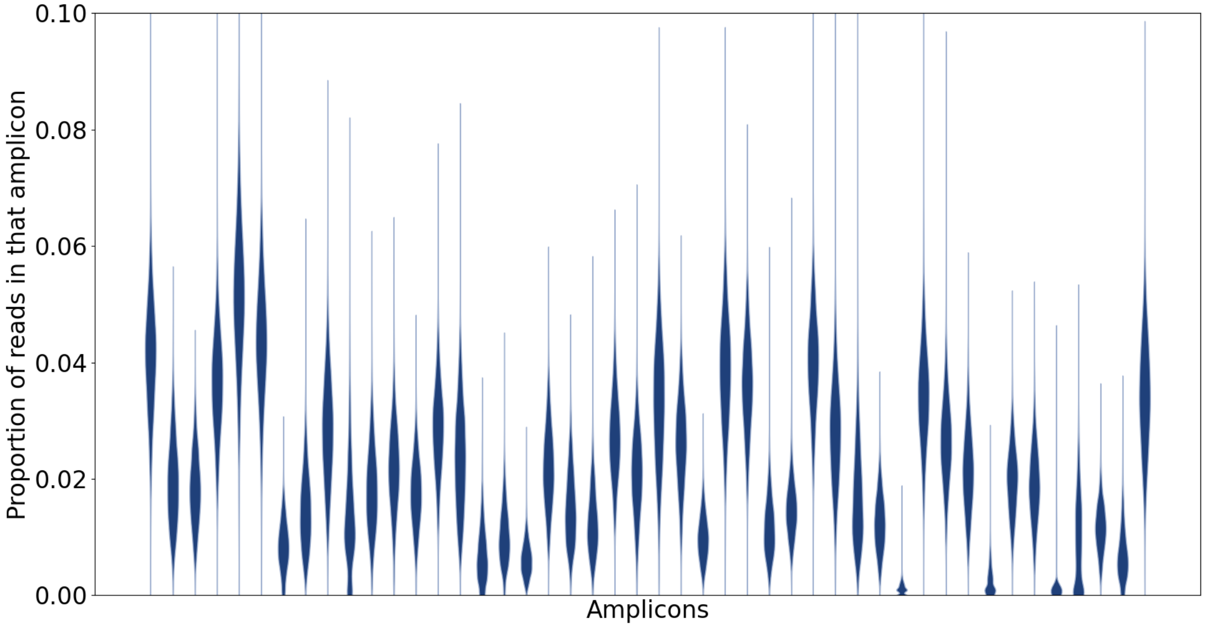


Supplementary Figure C.2: Violin pots of the proportion of reads falling on each amplicon for the sample AML-02-001 (50-amplicon panel). Each amplicon has a different coverage and, within one amplicon, there is a lot of variability across cells.

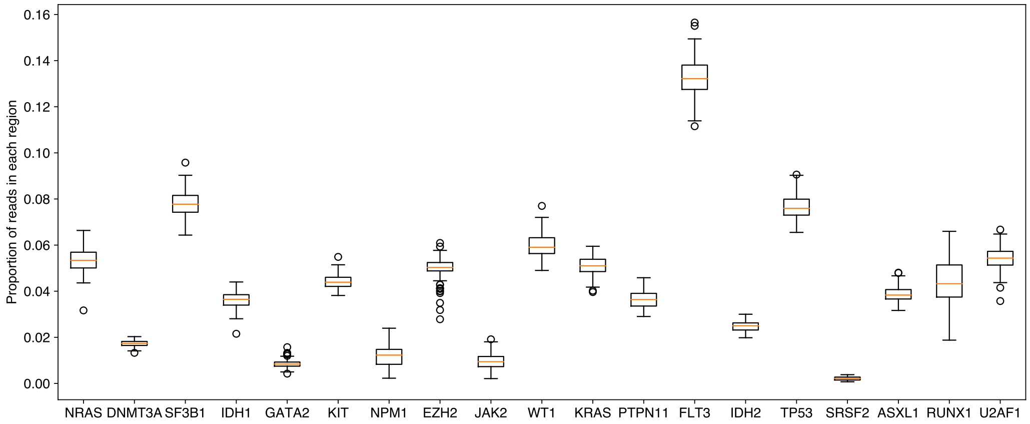


Supplementary Figure C.3: Boxplot of the median proportion of reads falling on each region in each sample, for samples sequenced with the 50-amplicon panel.

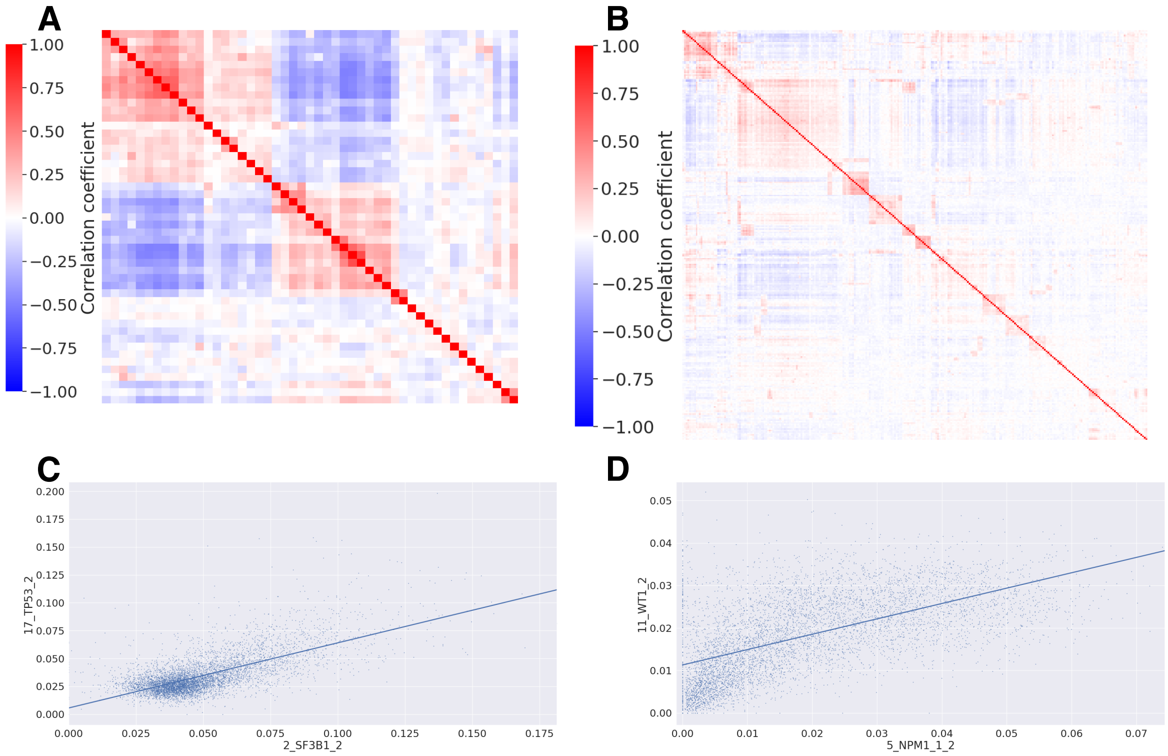


Supplementary Figure C.4: Correlations observed between the coverage of different amplicons in real MissionBio data. **A**. Correlation coefficients between the read count proportions on each amplicon in sample AML-08-001, sequenced with the 50-amplicon panel. The amplicons were sorted according to a hierarchical clustering. **B**. Correlation coefficients between the read count proportions on each amplicon in sample AML-89-001, sequenced with the 279-amplicon panel. **C**. Proportions of reads falling on two correlated amplicons in each cell: one amplicon on *TP53* (chr17) and one amplicon on *SF3B1* (chr2). **D**.:Proportions of reads falling on two correlated amplicons in each cell: one amplicon on *NPM1* (chr5) and one one *WT1* (chr11).

#### C.2 Allele dropout rates

Like every single-cell DNA sequencing platform, Tapestri® suffers from allelic dropouts: even if a cell is heterozygous, it can happen that only one of the two alleles is amplified, and the cell appears homozygous. The 50-amplicon panel used by Morita et al. contained 10 amplicons targeting common SNPs in order to estimate dropout rates. For each sample, we selected the SNPs for which it is heterozygous, and we computed the dropout rate of each allele: we define the dropout rate of the alternative allele as the number of cells genotyped as wild type divided by the total number of cells genotyped, and likewise for the dropout rate of the reference allele. The dropout rates of each allele are around 5%, but they vary a lot from variant to variant and from sample to sample (Supplementary Figure C.5). Apart from some outliers, both alleles have the same dropout rate.

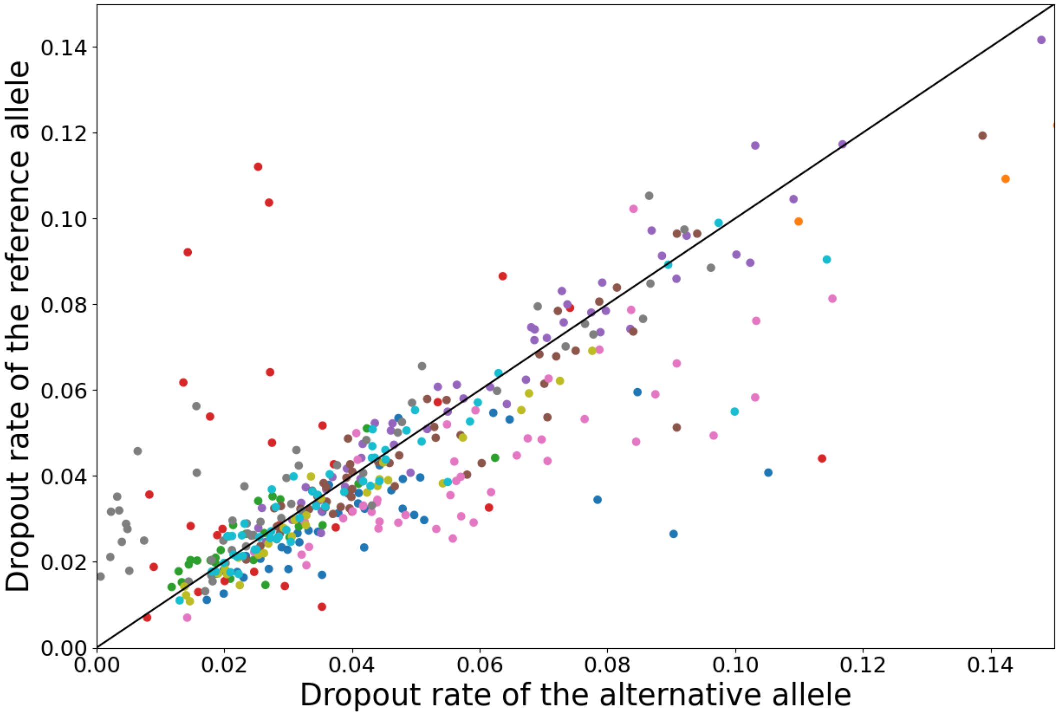


Supplementary Figure C.5: Dropout rates of the two alleles, for 10 common SNPs targeted by specific amplicons. Each point corresponds to a pair (sample, SNP), and points corresponding to the same SNP are grouped by colour. In most cases, both alleles have the same dropout rate, but the dropout rate varies a lot from variant to variant.

#### C.3 Allele frequencies

If one cell contains one copy of the wild type allele and one copy of the mutated allele, we expect in the absence of dropouts to find approximately 50% reads supporting each allele, with some stochastic variability. This is indeed what we observe in the Tapestri® data (Supplementary Figure C.6, top). In case of a copy number gain, there can be for example two copies of the reference allele and one copy of the alternative allele, in which case we would expect the alternative allele fraction to be centered on 1/3. In addition, it is very unlikely that both wild type copies fail to be amplified, which means that we don't expect to find many dropouts of the reference allele. This is indeed what is observed in real data (Supplementary Figure C.6, bottom left). For the larger panel with 279 amplicons, the alternative allele fraction distribution is not as peaked (Supplementary Figure C.6, bottom right), so in this case we use a larger overdispersion in the beta-binomial model.

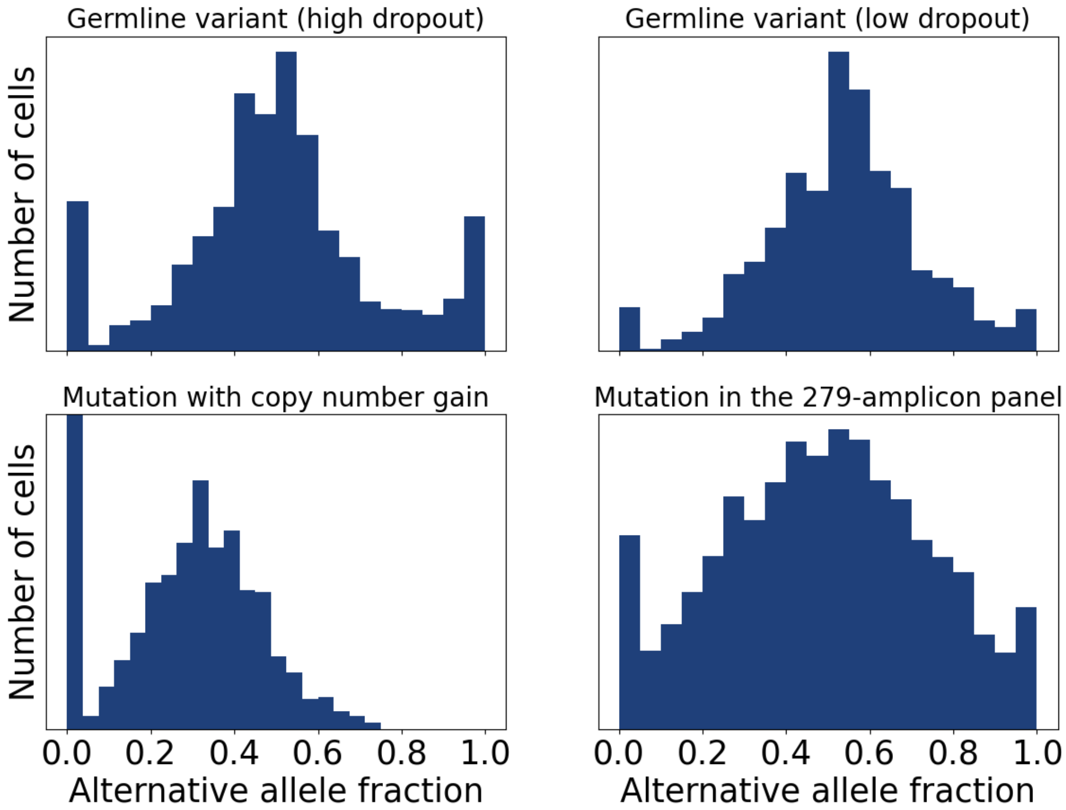


Supplementary Figure C.6: Fraction of alternative allele for two germline variants in sample AML-02-001 (top), *FLT3*-ITD in sample AML-28-001 (bottom left) and a mutation in *IDH1* in sample AML-82-001 (bottom right).

#### C.4 Genes covered by the panels

| Chrom arm | Gene | Nb amplicons in small panel | Nb amplicons in large panel |
|---|---|---|---|
| 1p | CSF3R | 0 | 4 |
| 1p | MPL | 0 | 1 |
| 1p | NRAS | 2 | 2 |
| 2p | DNMT3A | 1 | 22 |
| 2q | SF3B1 | 2 | 5 |
| 2q | IDH1 | 1 | 1 |
| 3q | GATA2 | 1 | 2 |
| 4q | KIT | 2 | 6 |
| 4q | TET2 | 0 | 26 |
| 5q | NPM1 | 1 | 1 |
| 7q | EZH2 | 2 | 19 |
| 8q | RAD21 | 0 | 4 |
| 8q | MYC | 0 | 6 |
| 9p | JAK2 | 1 | 2 |
| 10q | SMC3 | 0 | 4 |
| 11p | WT1 | 3 | 3 |
| 11q | CBL | 0 | 3 |
| 12p | ETV6 | 0 | 10 |
| 12p | KRAS | 2 | 3 |
| 12q | PTPN11 | 2 | 4 |
| 13q | FLT3 | 5 | 5 |
| 15q | IDH2 | 1 | 1 |
| 17p | TP53 | 4 | 9 |
| 17q | NF1 | 0 | 28 |
| 17q | PPM1D | 0 | 3 |
| 17q | SRSF2 | 1 | 1 |
| 18q | SETBP1 | 0 | 6 |
| 19p | CALR | 0 | 1 |
| 20q | ASXL1 | 2 | 10 |
| 21q | RUNX1 | 5 | 9 |
| 21q | U2AF1 | 2 | 2 |
| Xp | ZRSR2 | 0 | 11 |
| Xp | KDM6A | 0 | 19 |
| Xq | BCOR | 0 | 22 |
| Xq | SMC1A | 0 | 3 |
| Xq | STAG2 | 0 | 13 |
| Xq | PHF6 | 0 | 8 |
| 6,10,14,18 | SNPs | 10 | 0 |
| | Total | 50 | 279 |

Supplementary Table 1: List of genes targeted by each of the 2 panels used in the publication of Morita et al. (2020), with the chromosome arm in which they are located and the number of amplicons targeting them. The 50-amplicon panel contains 10 amplicons targeting common SNPs, which were used to infer allelic dropout rates. Genes targeted by only a few unreliable amplicons with a very high dropout rate were excluded from the CNA inference. For example, both panels had only one amplicon on 5q (targeting *NPM1*) and this amplicon was very unreliable and was therefore filtered out from the CNA inference, which is why we did not detect any deletion on 5q.

### D Simulations with synthetic data

#### D.1 Data generation

We generated synthetic data in a way that attempts to mimic real data, as is described below.

- The tree structure of $n_{\text{nodes}} = 6$ nodes is generated by randomly sampling a Prüfer sequence uniformly from $[\![0, n_{\text{nodes}} - 1]\!]^{n_{\text{nodes}}-2}$.
- $n_{\text{SNVs}}$ SNVs are randomly assigned to $n_{\text{regions}}$ regions.
- $n_{\text{SNVs}}$ SNVs, $n_{\text{CNAs}}$ CNAs are randomly assigned to the nodes in the tree, making sure that each node (except the root) contains at least one SNV or CNA.
- Each CNA is assigned randomly to one region. The regions are sampled without replacement. If the region contains a variant which is present in this node, randomly select the CNA type between gain, loss or CNLOH. Otherwise, select only between gain and loss. For each SNV present in the affected region (if any), we randomly select which allele is affected, making sure that the affected allele is present in this node.
- The node probabilities $\pi_1, ..., \pi_{n_{\text{nodes}}}$ (the probability that a cell is sampled from each node) are sampled from a uniform Dirichlet distribution whose concentration parameter is (10,...,10) in all simulations except the one where it is explicitly varied.
- For all variants, each allele has the same probability of 5% of being dropped out (except in the simulation where we varied the dropout rate).
- The region weights $\rho_k$ are sampled from a Normal distribution $N(1, \sigma)$ where $\sigma$ is 0.5 in all simulations except the one where it is explicitly varied. We set the minimum value of $\rho_k$ to be 0.3 to ensure that each region has a sufficient coverage.
- For each cell 3000 in total):
  - Sample the node to which it is attached from Categorical$(\pi_1, ..., \pi_{n_{\text{nodes}}})$.
  - For each variant $i$:
    * Sample the number of reads covering locus $i$ from a Poisson distribution with rate $20\rho_i$ (20 is the average sequencing depth).
    * Each allele copy can be dropped out with probability 5%. Let $f = \frac{c^{(a)}}{c^{(r)}+c^{(a)}}(1-\varepsilon) + \frac{c^{(r)}}{c^{(r)}+c^{(a)}}\varepsilon$ be the expected proportion of mutated reads, where $\varepsilon$ is the sequencing error rate which we set to 1% and $c^{(r)}$ and $c^{(a)}$ are the number of copies of the reference and alternative alleles which were not dropped out. Sample the number of mutated reads from a binomial distribution with parameter $f$, where the total number of reads is the one that was sampled from the Poisson distribution.

For Figure 2B, we used three CNAs per tree and removed CNLOH, in order to focus on the impact of the variance in region coverage. For Figure 2F, we used trees with 6 nodes and 5 SNVs (so one SNV per node, plus the root), since SCITE outputs trees with one SNV per node.

#### D.2 Parameters for the inference methods

BiTSC$^{\mathbf{2}}$ takes as input total and alternative read counts for each locus. Following the recommendation from BiTSC$^{\mathbf{2}}$'s developers, in cases where some regions do not contain any variants, we created one variant per region, where we set the alternative read count to 0. This results in all of these additional ”variants” to be placed by $\text{BiTSC}^2$ in one node where no cells are attached. Therefore, we post-processed $\text{BiTSC}^2$'s trees by removing the node with only placeholder SNVs. $\text{BiTSC}^2$ was run with 5000 burn-in samples and 500 actual samples. We ran 5 chains in parallel, using the default temperature parameter. $\text{BiTSC}^2$ normally performs the inference with different numbers of clones, and then selects the best model with a Bayesian Information Criterion. In order to speed up the simulations, we directly gave $\text{BiTSC}^2$ the correct number of clones (plus one, accounting for the node which we then remove in the post-processing), which also gives it an advantage. $\text{BiTSC}^2$ takes as input genomic segments, which divide the genome into groups of loci which share the same copy number. Here, we used regions as segments.

$\infty$SCITE outputs trees where one node contains only one SNV, whereas here nodes may contain several SNVs and CNAs. However, when one node in the true tree contains several SNVs, this will be represented in the SCITE tree by several successive nodes, where most of the intermediate nodes have no cells attached to them. Therefore, we post-processed SCITE's trees by deleting nodes with exactly one child and which have less than 2% of the cells attached to them, and moved their SNV to their child. This post-processing was not applied to the simulations where we varied the Dirichlet concentration parameter for the node probabilities, since in this case each node had exactly one SNV. SCITE was run with 4 different chains, each of length 20000. We gave SCITE the correct sequencing error and dropout rates, 1% and 5% respectively.

COMPASS was run with 4 parallel chains, each chain had 10000 iterations, and the temperature parameter was set to 10. Doublets were not modelled since the simulated data did not contain any.

#### D.3 Additional simulations

##### D.3.1 Number of cells for BiTSC²

In the simulations, we used 3000 cells, which is in the lower range of a typical Tapestri® dataset (1000 to 10000 cells). Surprisingly, we observed that BiTSC² performed worse when its input contained too many cells. Supplementary Figure D.1 shows that its performance dropped with an increasing number of cells. In contrast to COMPASS which marginalizes out the attachments of cells to nodes, BiTSC² samples the attachments of cells in the MCMC with a Gibbs sampling. This makes the inference much slower when there are many cells since many iterations are required to change the assignments of cells. In our simulation for Supplementary Figure D.1, we set the number of iterations for BiTSC² to 5000, which took more than 24 hours with trees of 6 nodes, 30 regions, 20 SNVs and 3 CNAs for 1000 cells. Despite this very long runtime, BiTSC² did not perform as well when given 500 or 1000 cells as input as with 100 or 200 cells. Consequently, it might be that the Gibbs sampling of the attachments of cells to nodes in BiTSC² makes the MCMC more likely to get stuck in a local optimum because, in order to reach a better tree, it might be necessary to change the assignments of a very large number of cells. This could explain the decreased performance of BiTSC² with a large number of cells. In all other simulations and for real data, we subsampled the input of BiTSC² to 200 cells, which improved its performance and decreased its runtime.

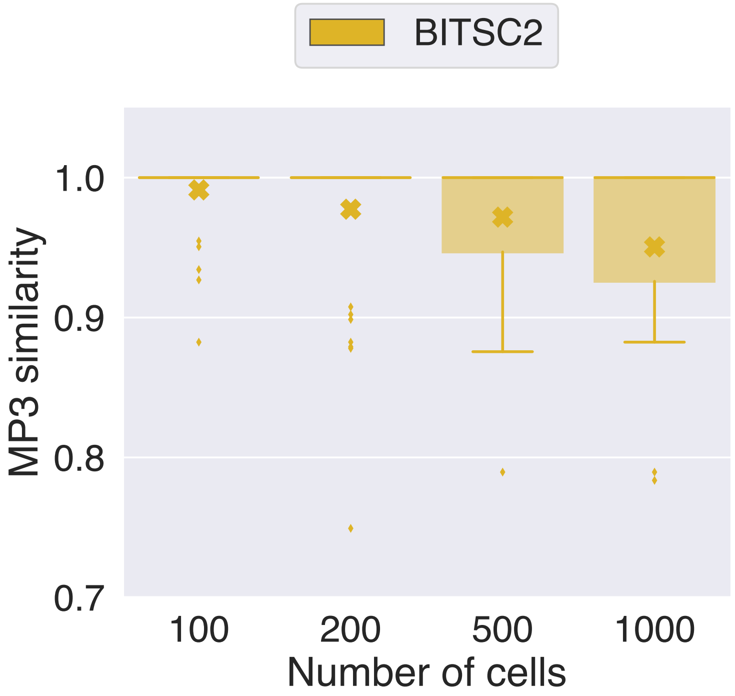


Supplementary Figure D.1: Evaluation of BiTSC² on synthetic data, with different numbers of cells. Here, we used trees with 6 nodes, 20 regions, 20 SNVs and 3 CNAs. For this simulation, we used the same coverage for all regions and disabled CNLOH. For each setting, we generated 50 different trees. The crosses represent the mean MP3 similarity to the true tree and the boxplots show the median and quartiles.

##### D.3.2 Doublets

In order to investigate the impact of doublets on the performance of the different methods, we also generated data with doublets, where we varied the doublet rate. COMPASS and ∞SCITE take doublets into account in their model, while BiTSC$^2$ does not. In order to focus on the impact of the doublets, we generated here data without CNAs.

Overall, all methods are very robust to doublets (Supplementary Figure D.2A). Only when the doublet rate is extremely high (90%, much higher than what would be found in real data) do we see a drop in performance for COMPASS and SCITE when they do not model doublets. When they account for doublets, their performance is almost not impacted by the presence of doublets in the data. Modelling doublets comes with an increased computational cost, which is more modest for COMPASS than SCITE (Supplementary Figure D.2B). Although both COMPASS and SCITE model doublets in a similar way, by considering pairs of nodes, SCITE is slower because the number of nodes in the trees is equal to the number of SNVs, whereas COMPASS can have several events in the same node, leading to a lower number of nodes. Since the runtime when modelling doublets is quadratic in the number of nodes, a lower number of nodes makes COMPASS much faster.

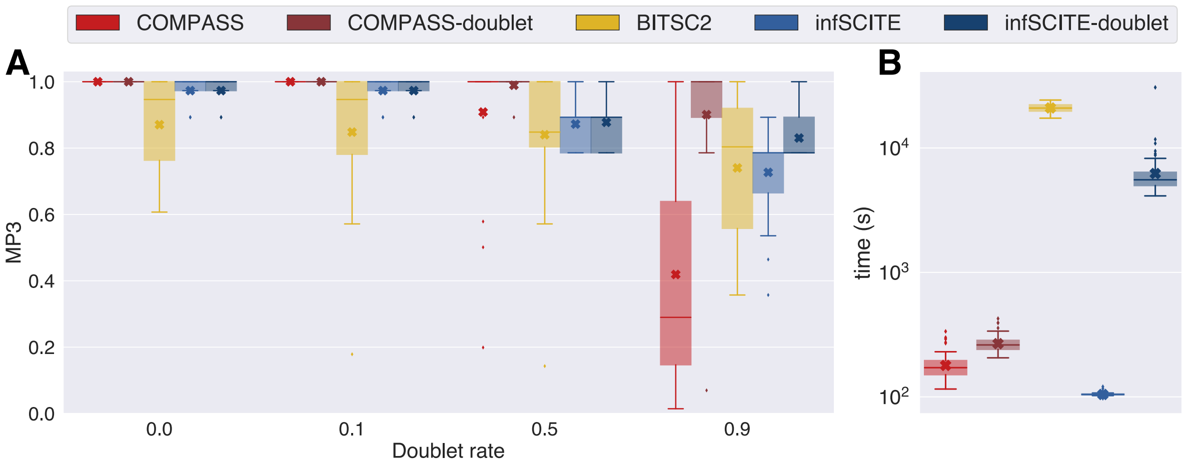


Supplementary Figure D.2: **A**. Evaluation of COMPASS, BiTSC$^2$ and SCITE on synthetic data, with different doublet rates. Here, we used trees with 6 nodes, 7 SNVs and 0 CNAs and a variance in region weights of 0. For each setting, we generated 50 different trees. The crosses represent the mean MP3 similarity to the true tree and the boxplots show the median and quartiles. **B**. Runtimes of COMPASS (with and without modelling doublets), BiTSC$^2$ and SCITE (with and without modelling doublets).

#### D.4 Correlations between regions

We generated data where regions were randomly clustered, and within each cluster the coverage on the regions were correlated, independently of CNAs, as we had observed to be the case in real data. Supplementary Figures D.3A-C show the correlations between regions with this synthetic data generation, which are similar the correlations observed in MissionBio's Tapestri® data. Supplementary Figure D.3D shows that the correlations of this type do not affect the results of COMPASS.

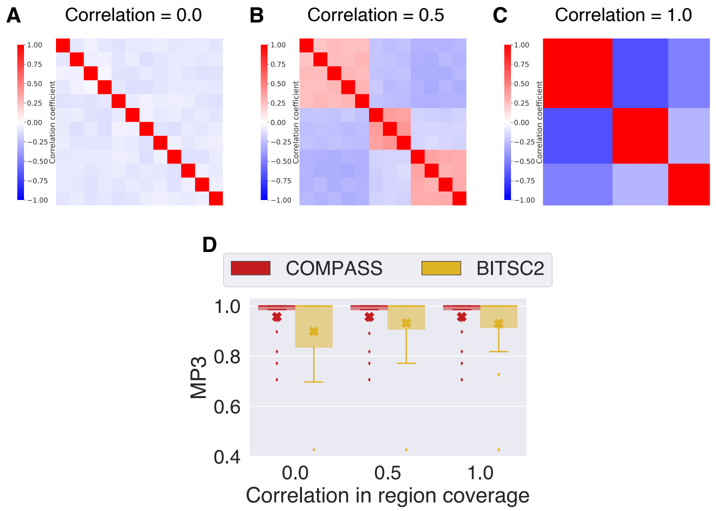


Supplementary Figure D.3: Evaluation of COMPASS and BiTSC² on synthetic data, with different values for the correlation coefficient of regions in the same cluster. The crosses represent the mean MP3 similarity to the true tree and the boxplots show the median and quartiles. The performances of COMPASS and BiTSC² are not affected by these correlations.

### E Orthogonal validation of COMPASS-derived CNA calls with bulk data

Bulk targeted sequencing data covering 297 genes was available for 94 samples, 9 of which were excluded because they were too noisy. We used CNVkit to detect CNAs (only copy number gains and losses, no CNLOH) in these samples. In addition, we also had bulk SNP array data for 40 samples, 8 of which were excluded because they were too noisy. We used ASCAT to detect CNAs (including CNLOH) for these samples. Since bulk data cannot be used to reliably infer subclonal CNAs, we only considered events detected by COMPASS in more than 50% of the cells. The complete list of CNAs detected by COMPASS and whether or not they could be validated in bulk data is available in Supplementary Table 2.

50-amplicon panel

| Patient | CNA | Bulk validation |
|---|---|---|
| AML-01 | CNLOH FLT3 (chr13) | unavailable |
| AML-02 | CNLOH JAK2 (chr9) | too subclonal |
| AML-03 | CNLOH FLT3 (chr13) | YES |
| AML-07 | Loss RUNX1 (chr21) | unavailable |
| | Loss U2AF1 (chr21) | unavailable |
| | Gain FLT3 (chr13) | unavailable |
| AML-19 | CNLOH FLT3 (chr13) | unavailable |
| AML-21 | CNLOH FLT3 (chr13) | unavailable |
| AML-25 | CNLOH RUNX1 (chr21) | YES |
| AML-32 | CNLOH RUNX1 (chr21) | YES |
| AML-38 | CNLOH FLT3 (chr13) | unavailable |
| AML-39 | Loss EZH2 (chr7) | YES |
| AML-41 | CNLOH WT1 (chr11) | unavailable |
| AML-43 | CNLOH WT1 (chr11) | unavailable |
| AML-49 | CNLOH FLT3 (chr13) | unavailable |
| AML-51 | CNLOH FLT3 (chr13) | unavailable |
| AML-59 | Loss TP53 (chr17) | too subclonal |
| | Loss EZH2 (chr7) | YES |
| | Gain WT1 (chr11) | YES |
| AML-60 | Gain ASXL1 (chr20) | NO |
| AML-65 | CNLOH FLT3 (chr13) | unavailable |
| AML-73 | Loss RUNX1 (chr21) | YES |
| AML-97 | CNLOH FLT3 (chr13) | unavailable |
| | CNLOH TP53 (chr17) | unavailable |

279-amplicon panel

| Patient | CNA | Bulk validation |
|---|---|---|
| AML-11 | CNLOH CBL (chr11) | YES |
| AML-42 | Loss TET2 (chr4) | YES |
| AML-71 | CNLOH DNMT3A (chr2) | NO |
| | CNLOH RUNX1 (chr21) | YES |
| AML-72 | CNLOH TET2 (chr4) | unavailable |
| AML-77 | CNLOH CBL (chr11) | unavailable |
| AML-78 | Loss EZH2 (chr7) | unavailable |
| | CNLOH TP53 (chr17) | unavailable |
| | Loss MYC (chr8) | unavailable |
| | Loss ETV6 (chr12) | unavailable |
| | Loss RAD21 (chr8) | unavailable |
| AML-79 | Loss TP53 (chr17) | unavailable |
| | Loss U2AF1 (chr21) | unavailable |
| | Gain U2AF1 (chr21) | unavailable |
| | Loss EZH2 (chr7) | unavailable |
| | Loss SETBP1 (chr18) | unavailable |
| | Gain RUNX1 (chr21) | unavailable |
| AML-83 | Gain MYC (chr8) | YES |
| | Gain RAD21 (chr8) | YES |
| AML-84 | CNLOH FLT3 (chr13) | unavailable |
| AML-88 | CNLOH FLT3 (chr13) | YES |
| AML-89 | CNLOH JAK2 (chr9) | unavailable |
| AML-91 | CNLOH TET2 (chr4) | YES |
| AML-92 | CNLOH JAK2 (chr9) | too subclonal |
| AML-98 | Loss EZH2 (chr7) | YES |
| AML-99 | CNLOH RUNX1 (chr21) | YES |
| | Gain MYC (chr8) | YES |
| | Gain RAD21 (chr8) | YES |
| AML-101 | Loss PPM1D (chr17) | unavailable |
| | Loss NF1 (chr17) | unavailable |
| | Loss EZH2 (chr7) | unavailable |
| | Loss TP53 (chr17) | unavailable |
| AML-103 | Loss TET2 (chr4) | unavailable |
| AML-107 | Loss FLT3 (chr13) | unavailable |
| | Loss EZH2 (chr7) | unavailable |
| | Loss ETV6 (chr12) | unavailable |
| AML-111 | Loss EZH2 (chr7) | unavailable |
| AML-112 | CNLOH CSF3R (chr1) | unavailable |
| | Gain KIT (chr4) | unavailable |
| | CNLOH DNMT3A (chr2) | unavailable |
| AML-113 | CNLOH FLT3 (chr13) | unavailable |
| AML-114 | CNLOH EZH2 (chr7) | unavailable |
| AML-117 | Loss TP53 (chr17) | unavailable |
| | Gain MYC (chr8) | unavailable |
| | Gain RAD21 (chr8) | unavailable |

Supplementary Table 2: List of CNAs detected in the AML cohort, for samples sequenced with the 50-amplicon and with the 279-amplicon panel. If SNP array or bulk targeted sequencing was available for the corresponding samples, we indicated whether the CNAs detected by COMPASS were also detected in the bulk data.

#### E.1 CNAs detected by COMPASS

Among the 16 patients for whom we detected CNAs, we also had bulk SNP array data for 2 of them (samples AML-59-001 and AML-99-001, shown in the main text) and bulk targeted sequencing (with a larger panel covering 297 genes) for 6 of them. Among these 8 samples, all of the CNAs present in a majority of the cells identified by COMPASS were also detected by bulk sequencing, except for one (amplification of *ASXL1* in sample AML-60-001, Supplementary Figure E.1).

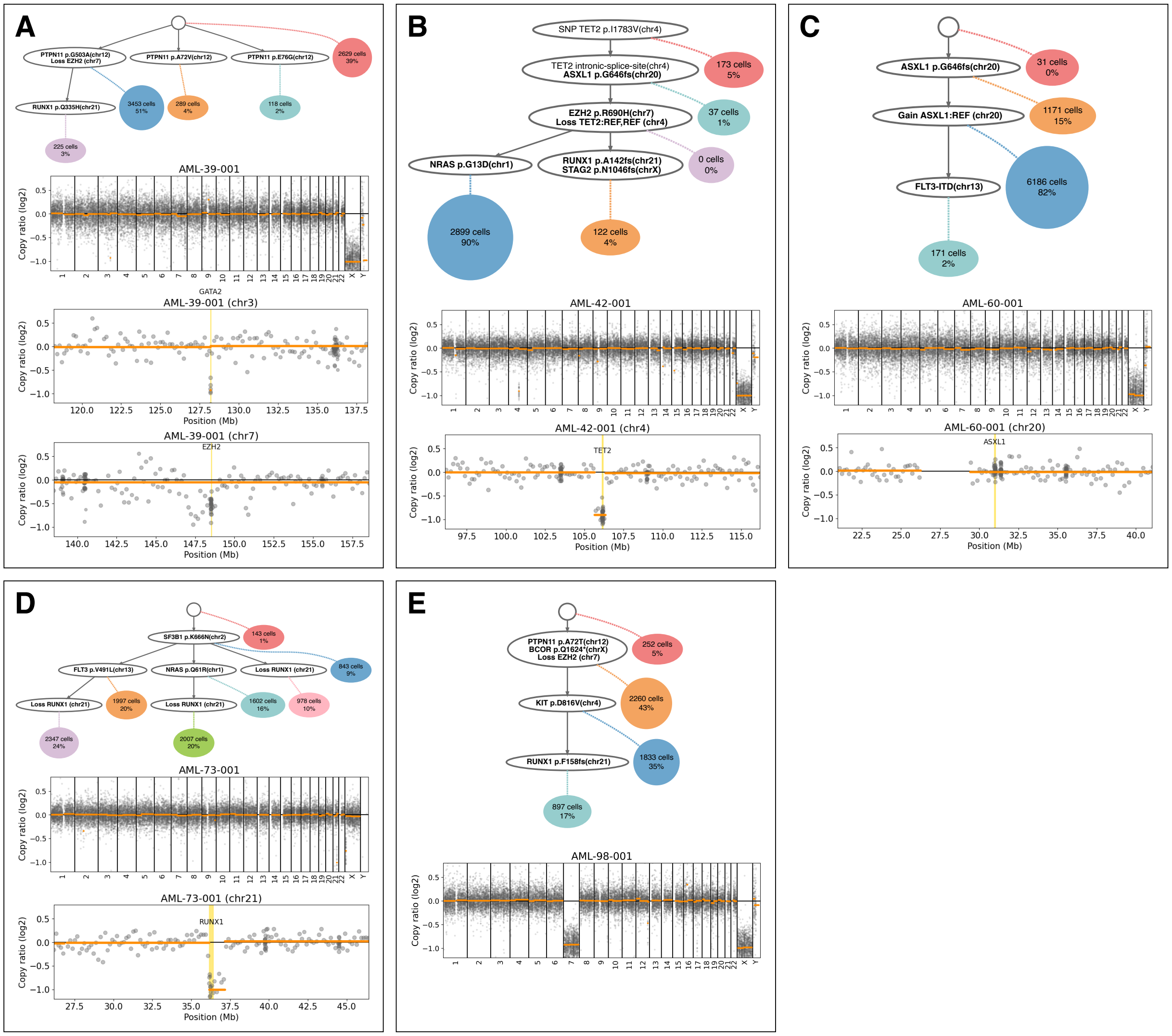


Supplementary Figure E.1: Trees inferred by COMPASS and corresponding copy-number plots generated by CNVkit from bulk sequencing for samples AML-39-001 (**A**), AML-42-001 (**B**), AML-60-001 (**C**), AML-73-001 (**D**), and AML-98-001 (**E**). CNVkit detected a small deletion of *GATA2* in sample AML-39-001 which was not detected by COMPASS because, since the amplicons on *GATA2* had a low coverage, this region was excluded from the CNA inference by COMPASS.

#### E.2 CNAs detected in bulk data but missed by COMPASS

Among the 85 samples for which we had reliable bulk targeted sequencing, some contained CNAs on regions not targeted by the single-cell panel, which obviously cannot be detected by COMPASS. Only one sample had CNAs that were identified by CNVkit in the bulk samples but were missed by COMPASS, even though they overlap regions targeted by the panel: AML-81-001 (chromosomes 8 and 9). This sample had a high number of cells attached to the root (47%). Since COMPASS estimates the region weights with the cells attached to the root, these weights may not be properly estimated when some neoplastic cells are attached to the root.

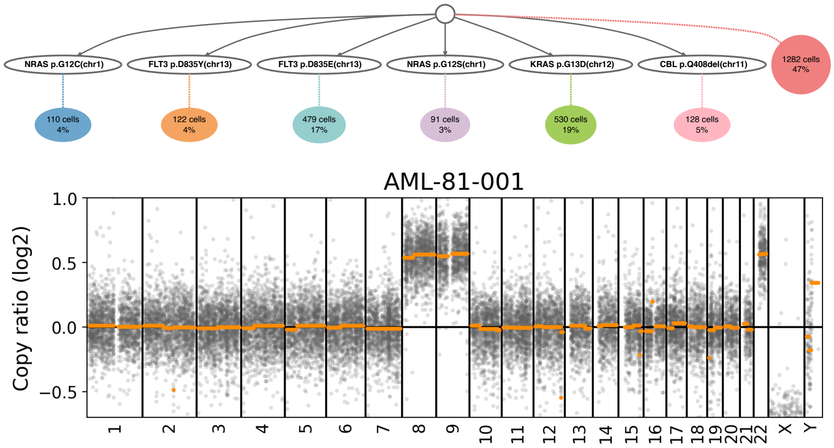


Supplementary Figure E.2: Tree inferred by COMPASS for sample AML-81-001, and copy-number profile inferred by CNVkit from bulk data. COMPASS did not identify any CNAs, even though CNVkit detected gains on chromosomes 8, 9 and 22. A high number of cells attached to the root (47%), which could have confounded the estimation of the region weights.

#### E.3 CNLOH events detected by COMPASS

Among the 32 samples for which reliable SNP array data was available, 6 of them contained CNLOH events detected by both COMPASS and ASCAT (AML-03-001, AML-11-001, AML-25-001, AML-71-001, AML-88-001, AML-91-001 and AML-99-001, Supplementary Figure E.3). Only in sample AML-71-001 did COMPASS detect a CNLOH event which was not validated by ASCAT (on *DNMT3A* on chr2). COMPASS assumes that allelic dropouts at different positions are independant, which may not be the case when several variants are on the same amplicon. Since this sample AML-71-001 contains two SNVs on *DNMT3A*, this assumption of the model might have led COMPASS to interpret the simultaneous dropouts of the two variants on *DNMT3A* as two CNLOH events.

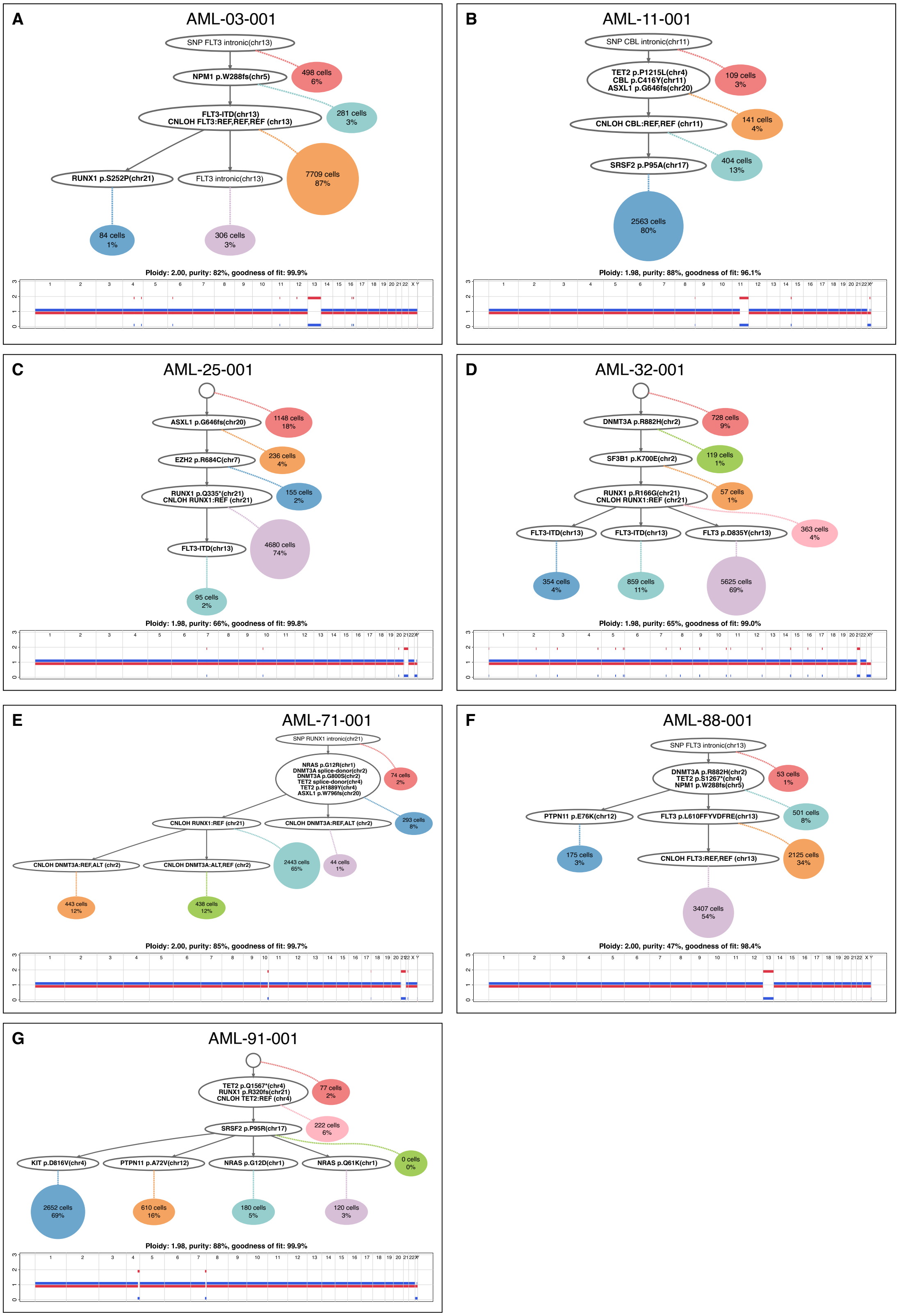


Supplementary Figure E.3: Trees inferred by COMPASS and corresponding ASCAT profiles for samples AML-03-001 (**A**), AML-11-01 (**B**), AML-25-001 (**C**), AML-32-001 (**D**), AML-71-001 ((**E**), AML-88-001 (**F**) and AML-91-001 (**G**).

##### E.4 CNLOH events detected with SNP array but missed by COMPASS

There are 4 samples for which we detected CNLOH events with SNP array data, but not with COMPASS: 19p in AML-20-001, 19q in AML-53-001, 10q in AML-71-001 and 7q in AML-91-001. All of these CNLOH events are either in regions not covered by the targeted panels or for which we did not detect SNVs, so there was no possibility for COMPASS to detect them.

### F Additional trees

#### F.1 Trees with CNAs

We show here all of the remaining trees where COMPASS inferred gains or losses, but for which we did not have bulk data to orthogonally validate the CNA calls.

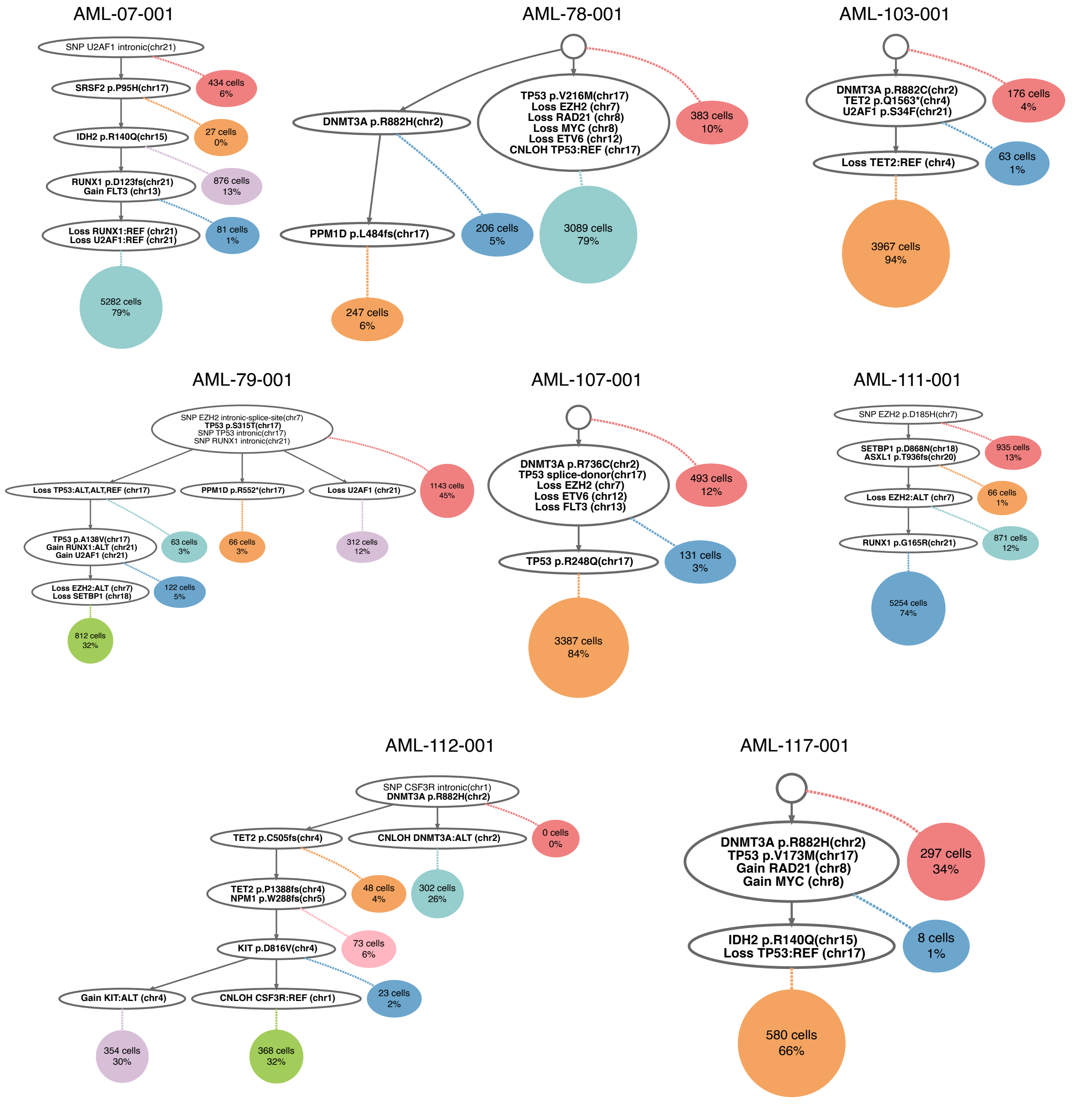


Supplementary Figure F.1: Inferred trees for samples for which COMPASS inferred gains or losses: AML-07-001, AML-78-001, AML-79-001, AML-103-001, AML-107-001, AML-111-001, AML-112-001, AML-117-001.

#### F.2 Longitudinal samples for patient AML-99

We had 5 longitudinal samples for patient AML-99, and the copy number gain on chromosome 8 and the CNLOH on *RUNX1* were detected in all of them.

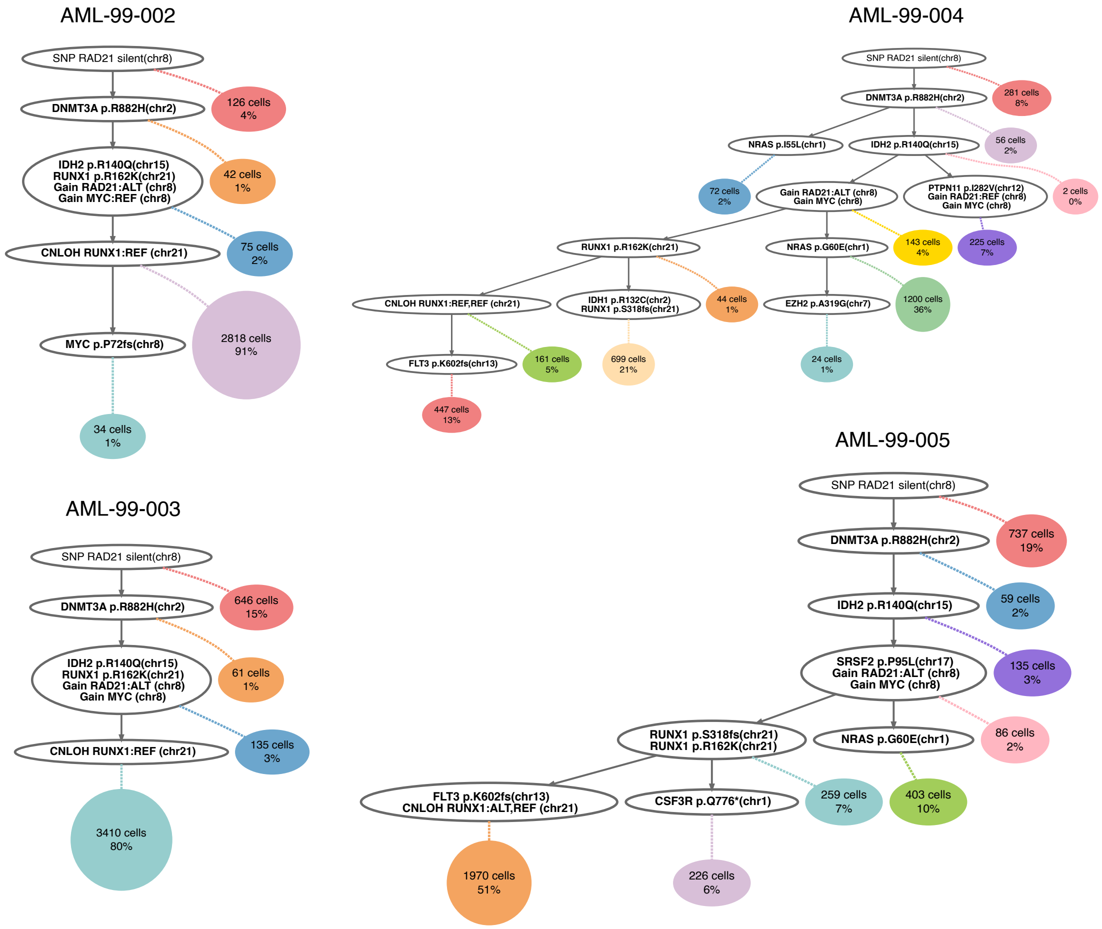


Supplementary Figure F.2: Inferred trees for the longitudinal samples of patient AML-99.

#### F.3 Trees with *JAK2* pV617F mutation and CNLOH

All 3 samples for which we detected the *JAK2* pV617F mutation also had a CNLOH at this locus.

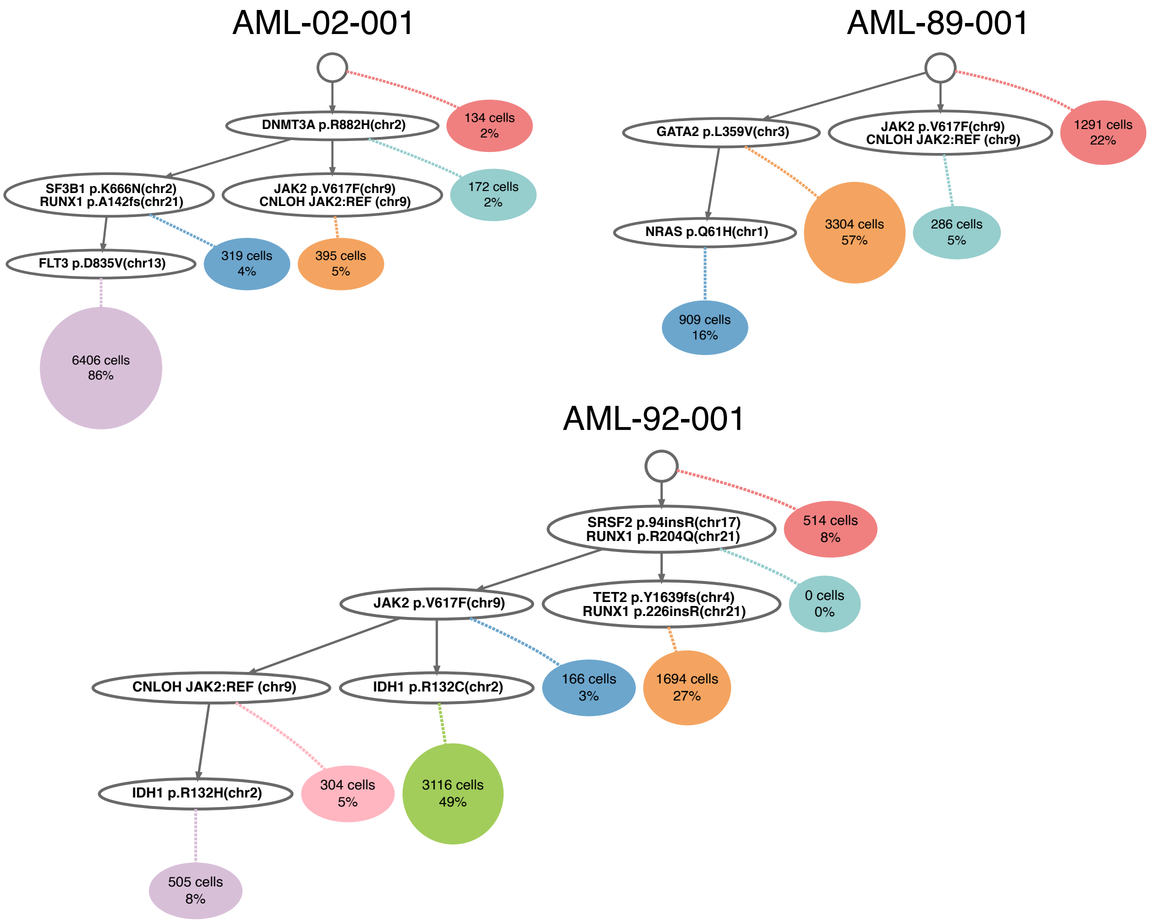


Supplementary Figure F.3: Inferred trees for the samples with *JAK2* mutations: AML-02-001, AML-89-001 and AML-92-001.

#### F.4 Samples with differences between COMPASS and SCITE

Since they are based on similar probabilistic models, COMPASS and ∞SCITE generally produce similar trees, at least when there are no CNAs. However, even in the absence of CNAs, there can be differences for two reasons.

First, when computing the likelihood of a tree, SCITE marginalizes over the attachments of cells to nodes, and assumes that each node has an equal prior probability, which is not a valid assumption when different clones have very different sizes. In such cases, SCITE might place a mutation corresponding to a small clone below a mutation corresponding to a larger clone, even when these two mutations actually occurred in parallel. The cells belonging to the larger clone can then also be attached to the node corresponding to the smaller subclone, albeit with a smaller likelihood since it requires a dropout of the mutation that they do not actually have. If the larger clone is much larger than the smaller one, the small gain in likelihood for a high number of cells can lead to a higher likelihood for the false tree. COMPASS allows each node to have a different attachment probability, which solves this problem. For example, in sample AML-02-001, COMPASS places the *JAK2* mutation in a different clone that the *SF3B1*, *RUNX1* and *FLT3* mutations, which is in accordance with the small number of cells which have both mutations and probably correspond to doublets (Supplementary Figure F.4A). However, SCITE places the *JAK2* mutation below the main clone, for the reasons explained above.

Another difference between SCITE and COMPASS is that SCITE assumes that all SNVs have the same dropout rate, whereas COMPASS allows each SNV to have its own dropout rate. This can lead to differences in the order of mutations. In general, if mutation A and mutation B are approximately present in the same cells but mutation A has a higher dropout rate, SCITE will place mutation A below mutation B in the tree. This is because there are more cells having mutation B without mutation A, than cells having mutation B without mutation A. For example, in sample AML-02-001, 175 cells have the *FLT3* mutation without the *RUNX1* mutation, whereas only 136 cells have the *RUNX1* mutation without the *FLT3* mutation (Supplementary Figure F.4A). Consequently, SCITE placed the *FLT3* mutation before *RUNX1*. However, among the *FLT3* mutated cells, there are as many *RUNX1* WT and mutated cells (175 and 165), indicating that, assuming both alleles are equally likely to be dropped out, all cells having the *RUNX1* mutation also have *FLT3*. Conversely, among cells with the *RUNX1* mutation, there are more cells without the *FLT3* mutation than with it (136 vs 84), indicating that the *FLT3* likely occurred after the *RUNX1* mutation, although the *RUNX1* mutation has a higher dropout rate. Similarly, in sample AML-16-001, the *DNMT3A* mutation has a higher dropout rate than *IDH1*, resulting in SCITE placing *DNMT3A* after *IDH1* (Supplementary Figure F.4B), even though the single-cell data dos not provide evidence for it, and *DNMT3A* mutations are known to be early events. A similar phenomenon is observed in sample AML-17-001 for *DNMT3A* and *NRAS* (Supplementary Figure F.4C).

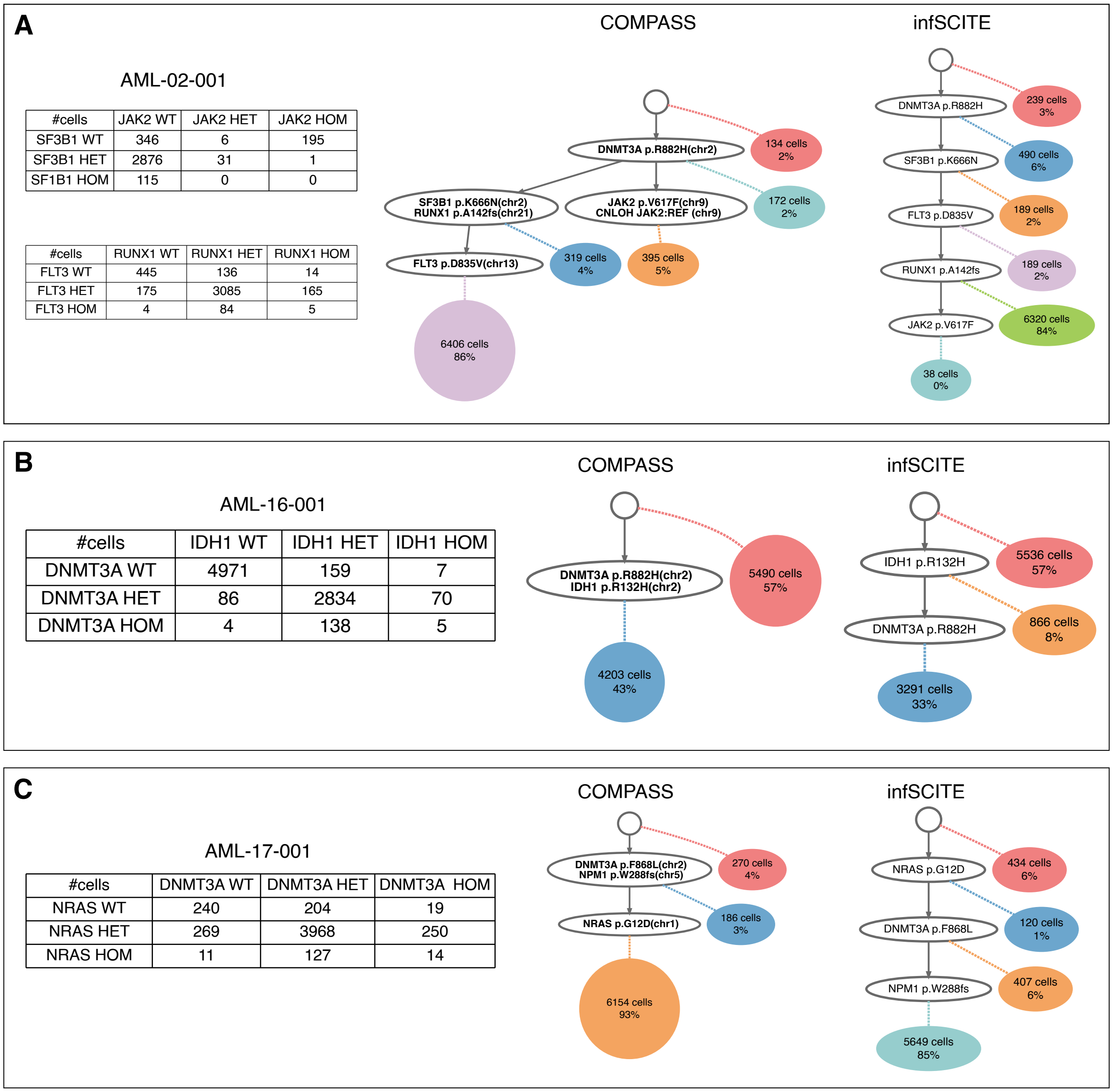


| #cells | JAK2 WT | JAK2 HET | JAK2 HOM |
|---|---|---|---|
| SF3B1 WT | 346 | 6 | 195 |
| SF3B1 HET | 2876 | 31 | 1 |
| SF1B1 HOM | 115 | 0 | 0 |

| #cells | RUNX1 WT | RUNX1 HET | RUNX1 HOM |
|---|---|---|---|
| FLT3 WT | 445 | 136 | 14 |
| FLT3 HET | 175 | 3085 | 165 |
| FLT3 HOM | 4 | 84 | 5 |

| #cells | IDH1 WT | IDH1 HET | IDH1 HOM |
|---|---|---|---|
| DNMT3A WT | 4971 | 159 | 7 |
| DNMT3A HET | 86 | 2834 | 70 |
| DNMT3A HOM | 4 | 138 | 5 |

| #cells | DNMT3A WT | DNMT3A HET | DNMT3A HOM |
|---|---|---|---|
| NRAS WT | 240 | 204 | 19 |
| NRAS HET | 269 | 3968 | 250 |
| NRAS HOM | 11 | 127 | 14 |

Supplementary Figure F.4: Example samples for which COMPASS and SCITE inferred a different order of the mutations. **A** Trees inferred by COMPASS and $\infty$SCITE for sample AML-02-001, and number of cells having each possible genotypes for the pairs of mutations (JAK2,SF3B1) and (RUNX1,FLT3). **B** Trees inferred by COMPASS and $\infty$SCITE for sample AML-16-001, and number of cells having each possible genotypes for the pair of mutations (DNMT3A,IDH1). **C** Trees inferred by COMPASS and $\infty$SCITE for sample AML-17-001, and number of cells having each possible genotypes for the pair of mutations (DNMT3A,NRAS).

#### F.5 Other AML samples with *TP53* mutations treated with venetoclax

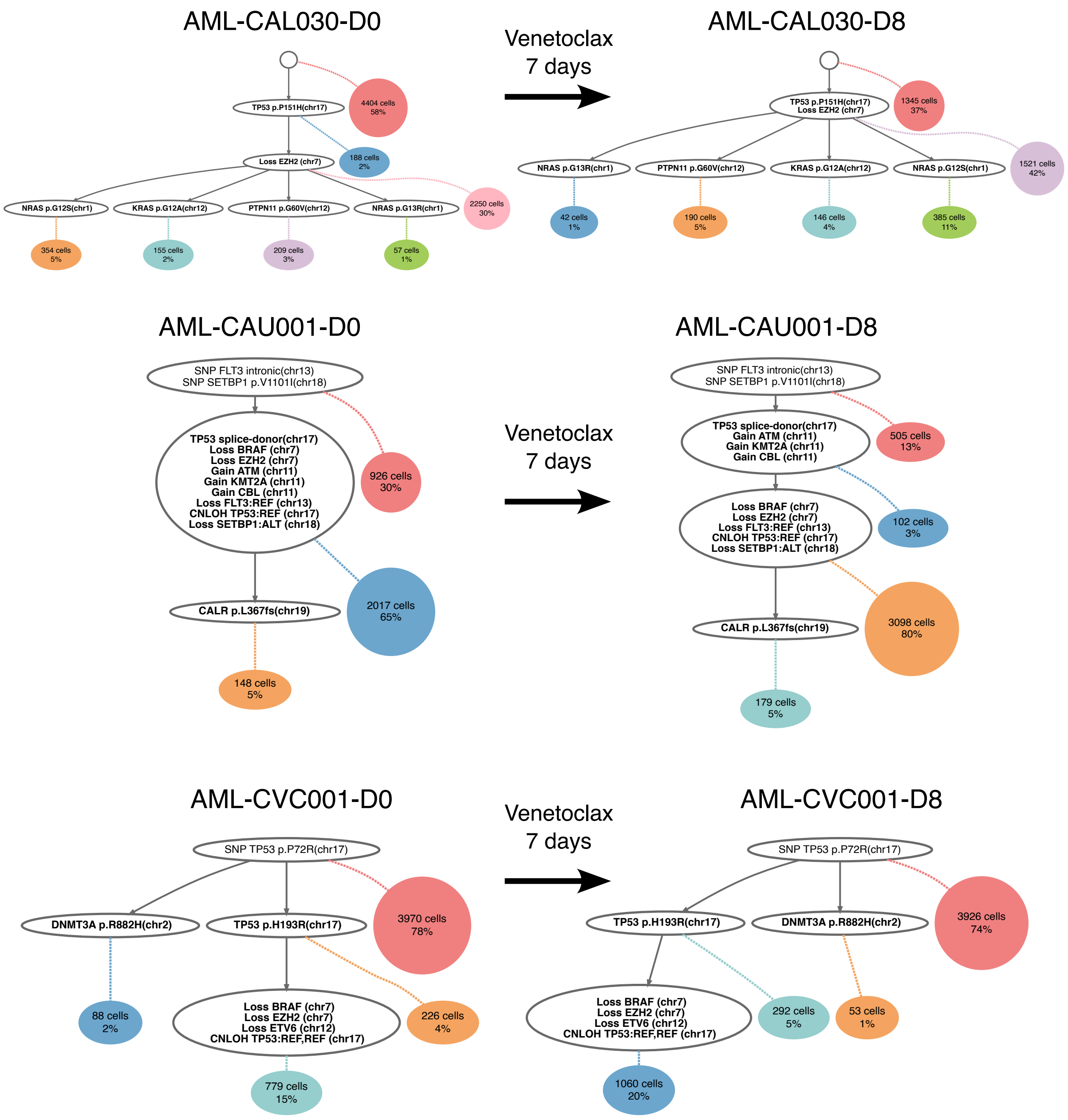


Supplementary Figure F.5: Inferred trees for the AML samples with *TP53* mutations treated with venetoclax, from Thijssen et al. 2021.

#### F.6 Other MPN samples with *TP53* mutations

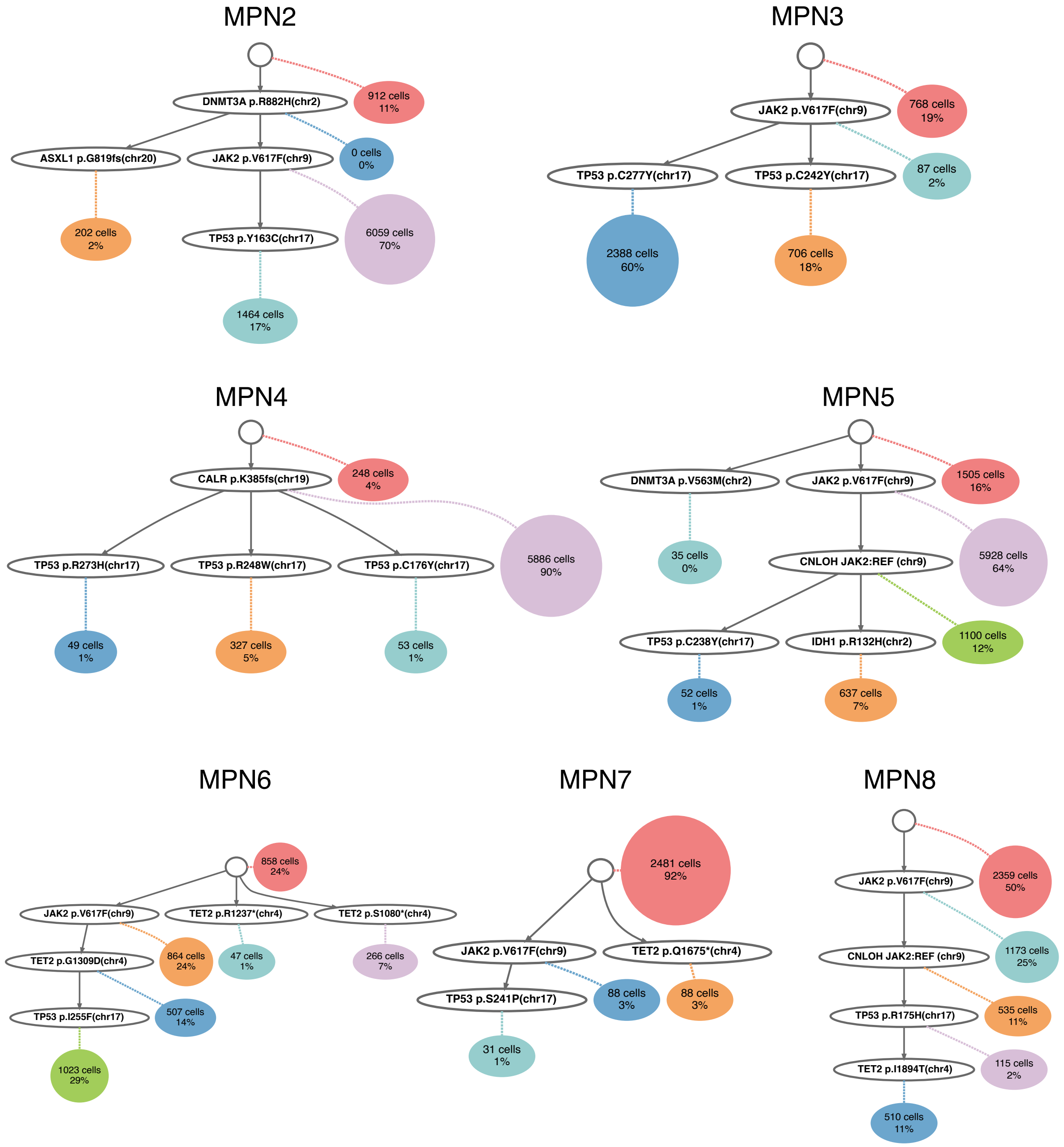


Supplementary Figure F.6: Inferred trees for the MPN samples with *TP53* mutations, from Maslah et al. 2022.
